# DDX50 is a viral restriction factor that enhances TRIF-dependent IRF3 activation

**DOI:** 10.1101/2020.08.06.239376

**Authors:** Mitchell A. Pallett, Yongxu Lu, Geoffrey L. Smith

## Abstract

The transcription factors IRF3 and NF-κB are crucial in innate immune signalling in response to many viral and bacterial pathogens. However, mechanisms leading to their activation remain incompletely understood. Canonical RLR signalling and detection of viral RNA is dependent upon the receptors RIG-I, MDA5 and TLR3. Alternatively, the DExD-Box RNA helicases DDX1-DDX21-DHX36 activate IRF3/NF-κB in a TRIF-dependent manner independent of RIG-I, MDA5 or TLR3. Here we describe DDX50, which shares 55.6% amino acid identity with DDX21, as a component of the dsRNA sensing machinery and signalling pathway. Deletion of DDX50 in mouse and human cells impaired activation of the IFNβ promoter, IRF3-dependent endogenous gene expression and cytokine/chemokine production in response to cytoplasmic dsRNA (polyIC transfection), and infection by RNA and DNA viruses. Mechanistically, DDX50 co-immunoprecipitated with TRIF and DDX1, promoting complex formation upon stimulation. Furthermore, whilst MAVs/TBK1 induced signalling is intact in *Ddx50* KO cells, TRIF-dependent signalling was impaired suggesting DDX50 drives TRIF-dependent *Ifnβ* transcription. Importantly, loss of DDX50 resulted in increased replication and dissemination of vaccinia virus, herpes simplex virus and Zika virus highlighting its important role as a viral restriction factor.

**Author summary:** The detection of viral RNA or DNA by host RNA or DNA sensors and the subsequent antiviral immune response are crucial for the outcome of infection and host survival in response to a multitude of viral pathogens. Detection of viral RNA or DNA culminates in the upregulation of inflammatory cytokines, chemokines and pathogen restriction factors that augment the host innate immune response, restrict viral replication and clear infection. The canonical RNA sensor RIG-I is a member of the large family of DExD/H-box helicases, however the biological role of many DExD/H-box helicases remain unknown. In this report, we describe the DExD-Box helicase DDX50 as a new component of the RNA sensing machinery. In response to DNA and RNA virus infection, DDX50 functions to enhance activation of the transcription factor IRF3, which enhances antiviral signalling. The biological importance of DDX50 is illustrated by its ability to restrict the establishment of viral infection and to diminish the yields of vaccinia virus, herpes simplex virus and Zika virus. These findings increase knowledge of the poorly characterised host protein DDX50 and add another factor to the intricate network of proteins involved in regulating antiviral signalling in response to infection.

## Introduction

Interferon regulatory factor 3 (IRF3)-dependent signalling leading to type I interferon (IFN) expression is crucial for pathogen clearance and host survival in response to infection by many viral and bacterial pathogens (1,2). IRF3 signalling is tightly regulated and is triggered by intracellular cytoplasmic/endoplasmic detection of viral RNA (dsRNA/5’-ppp/pp-RNA) (3) and DNA by pattern recognition receptors (PRRs) (4). The retinoic acid-inducible gene I (RIG-I)-like receptors (RLRs), which include DDX58 (RIG-I), DDX21, DDX1, DHX36, DDX60, DDX3, DHX9, DHX33, MDA5, and LPG2, as well as Toll-like receptor (TLR) −3, bind directly to, or form complexes with, viral dsRNA or 5’-ppp-RNA to activate downstream kinases and induce expression of type I IFN (5). RIG-I/MDA5 activation leads to mitochondrial antiviral signalling protein (MAVS)-dependent autophosphorylation of TANK-binding protein-1 (TBK1). In turn, TBK1 phosphorylates IRF3, leading to its dimerisation and nuclear translocation. In parallel, the transcription factor nuclear factor kappa-light-chain-enhancer of activated B cells (NF-κB) is activated in an inhibitor of nuclear factor kappa-B kinase subunit beta (IκKβ) dependent-manner (6). IRF3 and NF-κB trigger co-transcriptional upregulation of IFNs and inflammatory cytokines and chemokines, including IFNβ and C-X-C motif chemokine 10 (CXCL10/IP-10), as well as IRF3-dependent viral restriction factors (7,8). This establishes the host antiviral innate immune response, restricting viral replication and clearing infection.

The prototypical RLR, RIG-I, comprises a DExD-box ATPase-dependent RNA helicase with an N-terminal caspase activation and recruitment domain (CARD) and a C-terminal auto-inhibitory regulatory domain (RD). Briefly, under resting conditions, RIG-I is held in an autoinhibitory conformation. Upon agonist (5’-ppp/pp-RNA or short dsRNA) binding to the RD, RIG-I undergoes conformational change, dimerisation and activation (9). Through interaction with the E3 ligase tripartite-motif containing protein 25 (TRIM25) and subsequent K63-linked ubiquitylation of RIG-I, signalling is transduced via complex formation with the adaptor MAVS (10). Similarly, MDA5 signalling converges at MAVS activation, but differs from RIG-I receptor signalling due to alterations in ligand specificity. Alternatively, TLR3 differs in cellular compartmentalisation/localisation and signals in a TIR domain-containing adapter molecule 1 (TICAM-1 or TRIF) and TBK-dependent manner independent of MAVS. MDA5 recognises high molecular weight dsRNA (11) or mRNA lacking 2’-O-methylation at the 5’ cap (12), whereas TLR3 detects dsRNA in the endosomal or extracellular compartments. These subtle differences mean that during infection numerous RLRs and RNA sensors are activated in parallel or independent of one another and this is dependent upon cell type, the pathogen and/or the specific ligands present.

Other DExD-Box RNA helicase family members play an essential role in IRF3 signalling in response to viral PAMPs, including DDX60 (13,14), DDX1, DHX36, DDX21 (15), DHX33 (16) and DDX41 (17). Miyashita and colleagues identified DDX60 as a component of RLR-dependent signalling, acting through RIG-I and MDA5 to trigger optimal IRF3-dependent gene expression (13,14). However, a role for DDX60 in RLR signalling is contentious and recently it was reported to be dispensable for IFN production in response to several RLR agonists (18). Additionally, DDX1, DHX36 and DDX21 form a cytoplasmic complex with TRIF upon detection of 5’-ppp-RNA or dsRNA (PolyIC). Upon complex formation, DDX1 and DHX36 interaction, and TRIF recruitment, are DDX21-dependent, whereas DDX1 acts as the complex RNA sensor. Interestingly, this complex acts independently of TLR3, RIG-I and MDA5 in mouse dendritic cells (DCs) and mouse embryonic fibroblasts (MEFs) (15).

A recent RNAi screen, implicated the relatively uncharacterised DExD-Box RNA helicase proteins DDX17 and DDX50 as putative positive regulators of IFNβ promoter activity in response to cytoplasmic 5’-ppp-RNA (19). DDX50 is a paralogue of DDX21 sharing 55.6 % amino acid identity (20). DDX21 (Guα; nucleolar protein 2) and DDX50 (Guβ; nucleolar protein 1) are the only members of the Gu family of nucleolar RNA helicases and contain a highly homologous GUCT (Gu C-terminal) domain, which is followed by an arginine-serine-rich C-terminal tail in DDX50 (21). DDX50, as the name suggests, is localised to the nucleoli and *in vitro* assays have demonstrated that both DDX21 and DDX50 have ATPase and helicase activity, however DDX50 lacks RNA folding activity (21). Although DDX21 and DDX50 may have arisen by gene duplication on chromosome 10, these proteins have non-redundant roles. DDX21 targets RNA substrates with a 21- or 34-nt duplex and 5’-overhangs, whereas DDX50 targets only 21-nt duplex RNA for unwinding (21). On the other hand, DDX50 is required for optimal DDX21 unwinding activity, suggesting some co-dependence (21). Little is known about the biological function of DDX50, with one study suggesting it may be involved in MAP-kinase signalling through interaction with c-Jun (22).

By using CRISPR technology to knockout *Ddx50* in MEFs and *DDX50* in human embryonic kidney 293T cells (HEK293Ts), DDX50 was identified as a restriction factor for the DNA viruses vaccinia virus (VACV) and herpes simplex virus type 1 (HSV-1) and the RNA virus Zika (ZIKV). Mechanistically, DDX50 enhances IRF3-dependent gene transcription and cytokine synthesis and secretion in response to cytoplasmic dsRNA, and RNA or DNA virus infection.

## Results

### DDX50 is a novel factor required for nucleic acid sensing

To investigate the putative role of DDX50 in cytoplasmic RNA sensing, CRISPR-mediated knockouts (KO) were generated in MEFs and HEK293Ts. Successful KO was confirmed by immunoblotting (Fig. S1B and E) and genomic sequencing of individual alleles (Fig. S1A, C, D, F). Sequencing indicated frameshifts in exon 1 (Fig. S1C) and exon 4 (Fig. S1F) producing nonsense mutations and introduction of an early stop codon. No differences in morphology or growth properties between the wild type (WT) and KO cells were observed (data not shown). Initially, the contribution of DDX50 to IRF3 signalling in response to RLR agonists was investigated. Cells were co-transfected with a Firefly Luciferase reporter plasmid under the control of the IFNβ promoter (pIFNβ-Luc) and an internal control plasmid constitutively expressing Renilla Luciferase (pTK-RL). These cells were further mock-transfected or transfected with PolyIC (dsRNA analogue), infected with an RNA virus (Sendai virus (SeV)) or treated with extracellular PolyIC. IFNβ promoter activity was then measured relative to Renilla luminescence and non-stimulated controls. Promoter activity was significantly diminished in KO cells in comparison to WT cells in response to all stimuli (Fig. 1A), validating the results observed in the initial RNAi screen (19). Consistent with this observation, knockout of *Ddx50* reduced the expression of endogenous NF-κB and/or IRF3-dependent genes (*Isg56*, *Cxcl10* and *Ifnb*) in response to PolyIC and SeV as measured by RT-qPCR (Fig. 1B) and ELISA (CXCL10 and IL-6; Fig. 1C-D). Collectively, this indicates that DDX50 affects both the IRF3 or IRF3 and NF-κB branch of signalling in response to cytoplasmic dsRNA. Importantly, the defect in NF-κB/IRF3-dependent gene expression in response to PolyIC was rescued by transduction and complementation of *Ddx50* KO cells with a lentiviral vector encoding *Ddx50* (Fig. 2A-C). This ruled out CRISPR off target effects for the observed defect. Furthermore, overexpression of DDX50 augmented IFNβ promoter activity (Fig. S2A-B) and secretion of CXCL10 and IL-6 in response to PolyIC transfection (Fig. S2C-D). Interestingly, overexpression of DDX50 alone induced secretion of CXCL10 (Fig. S2C), indicating pathway activation above basal level even in the absence of stimulation. Concomitant experiments in HEK293T *DDX50* KO lines confirmed a dependence on DDX50 for optimal pathway activation in response to SeV infection in a human cell line (Fig. 1E-F).

**Fig. 1.**
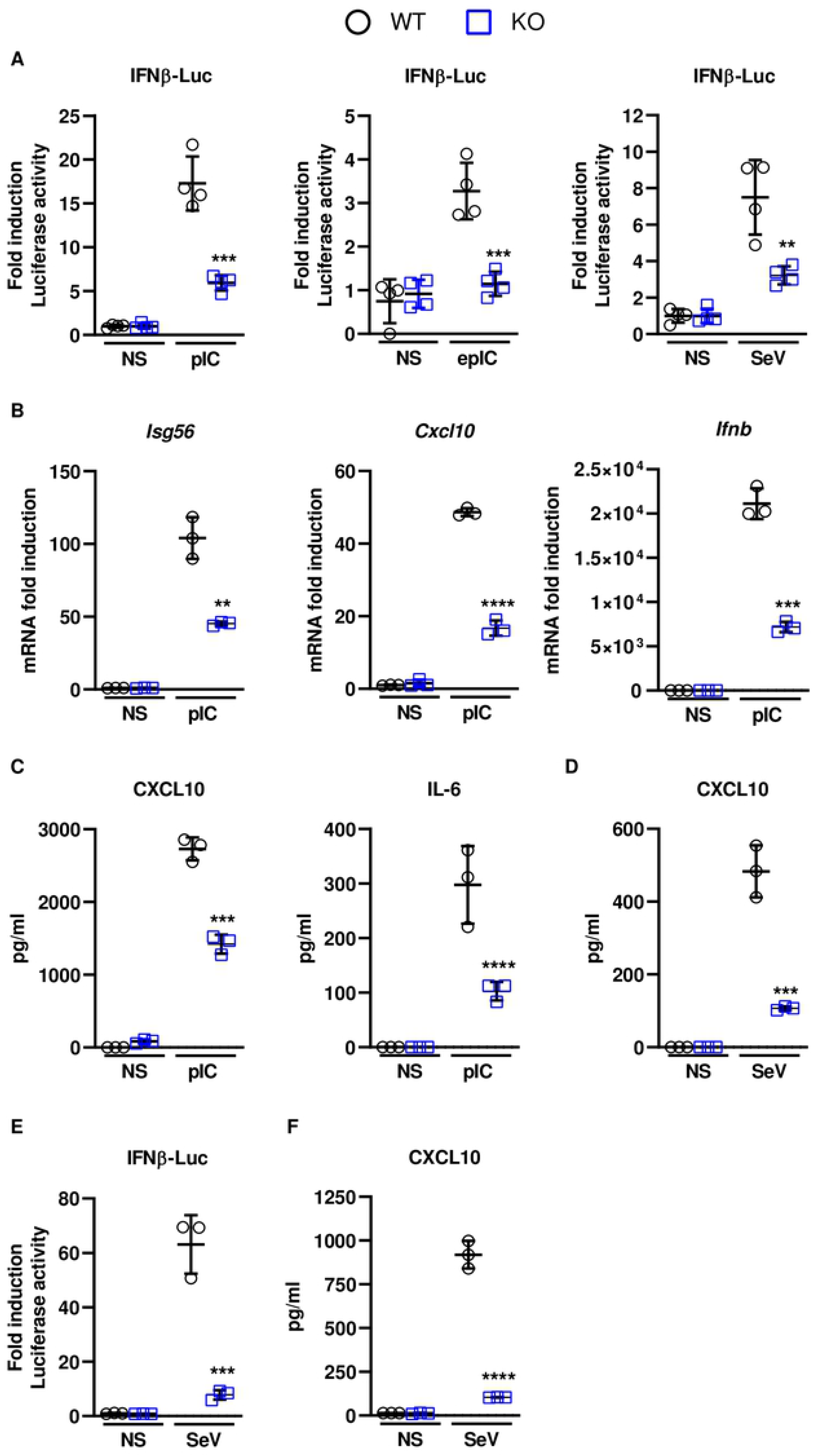
DDX50 (RH-II/Guβ) is required for intracellular nucleic acid sensing. Firefly luciferase activity of WT or *Ddx50*^*−/−*^ MEFs transfected with plasmids encoding Firefly Luciferase under the *Ifnβ* promoter and Renilla. **(A)** Cells were left untreated or treated with 5 μg/ml extracellular PolyIC (epIC), transfected with 5 μg/ml PolyIC (pIC) for 6 h or infected with Sendai virus (SeV) for 24 h. **(B)** WT or *Ddx50*^*−/−*^ MEFs were transfected with lipofectamine only or 5 μg/ml pIC for 7 h, and the fold induction of *Isg56*, *Cxcl10* or *Ifnb* mRNA levels, relative to *Gapdh,* were analysed by RT-qPCR. **(C)** Secreted levels of CXCL10 and IL-6 in the medium at 7 h post transfection with PolyIC or **(D)**4.5 h post infection with SeV were analysed by ELISA. **(E)** Firefly Luciferase activity of WT or *Ddx50*^*−/−*^ HEK293Ts transfected with plasmids encoding Firefly Luciferase under the *Ifnβ* promoter and Renilla. Cells were infected for 24 h with SeV or left untreated. Data are representative of at least three independent experiments. **(F)** Secreted levels of CXCL10 in the medium at 24 h post infection of WT or *Ddx50*^*−/−*^ HEK293Ts with SeV, were analysed by ELISA. Data are representative of at least three independent experiments. Statistical significance shown for WT stimulated vs KO stimulated.

**Fig. 2.**
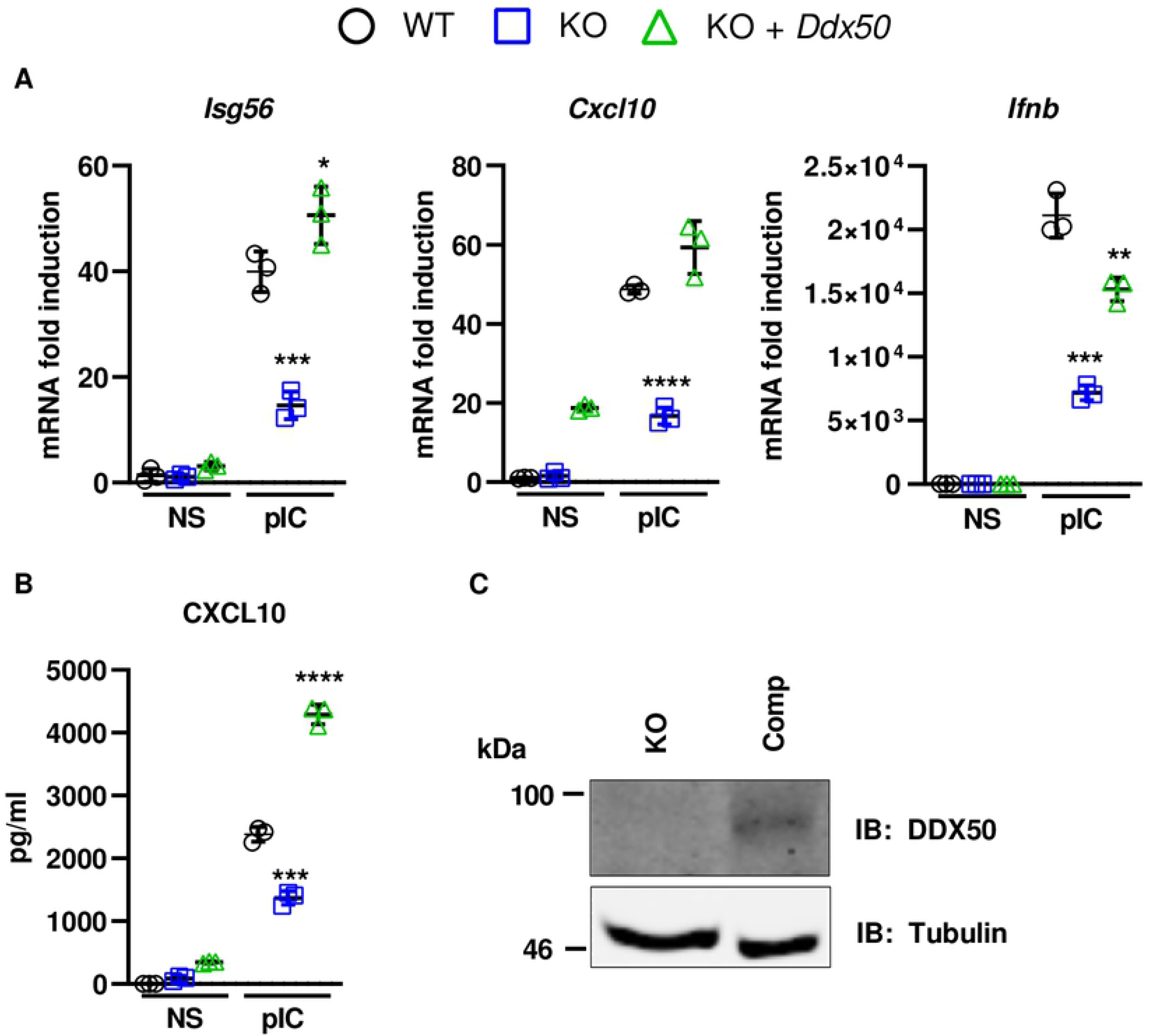
DDX50 rescues nucleic acid sensing in *Ddx50^−/−^* MEFs. **(A-C)** WT MEFs transduced with pLDT-EV and *Ddx50*^*−/−*^ MEFs transduced with pLDT-EV or pLDT-*Ddx50* were transfected with lipofectamine only or 5 μg/ml PolyIC for 7 h. **(A)***Isg56*, *Cxcl10* or *Ifnb* mRNA levels, relative to *Gapdh,* were analysed by RT-qPCR and **(B)** secreted CXCL10 was measured by ELISA. Representative of at least two independent experiments. Statistical significance shown for WT stimulated vs KO stimulated. **(C)** Expression of DDX50 was confirmed by SDS-PAGE and immunoblotting.

### DDX50 is required for IRF3/NF-κB driven gene expression during DNA virus infection

To investigate the biological relevance of DDX50, WT or KO MEFs were infected with two large dsDNA viruses, VACV and HSV-1. The modified vaccinia Ankara (MVA) strain and the HSV-1 ΔICPO strain each elicit strong innate immune responses in tissue culture and were therefore used to increase pathway activation and sensitivity, as described previously (23,24). Following infection with either MVA or HSV-1 ΔICP0, the expression of IRF3 and/or NF-κB-dependent *Isg56* and *Ifnb* were significantly diminished in the KO cells in comparison to infected WT cells (Fig. 3A-B). This correlated with decreased secretion of CXCL10 and IL-6 as determined by ELISA (Fig. 3C-D). Although significant, the effect of *Ddx50* KO on *Cxcl10* expression by RT-qPCR following infection with both viruses was less pronounced (Fig. 3A-B). Overall, this highlights the importance of DDX50 in innate immune signalling during viral infection.

**Fig. 3.**
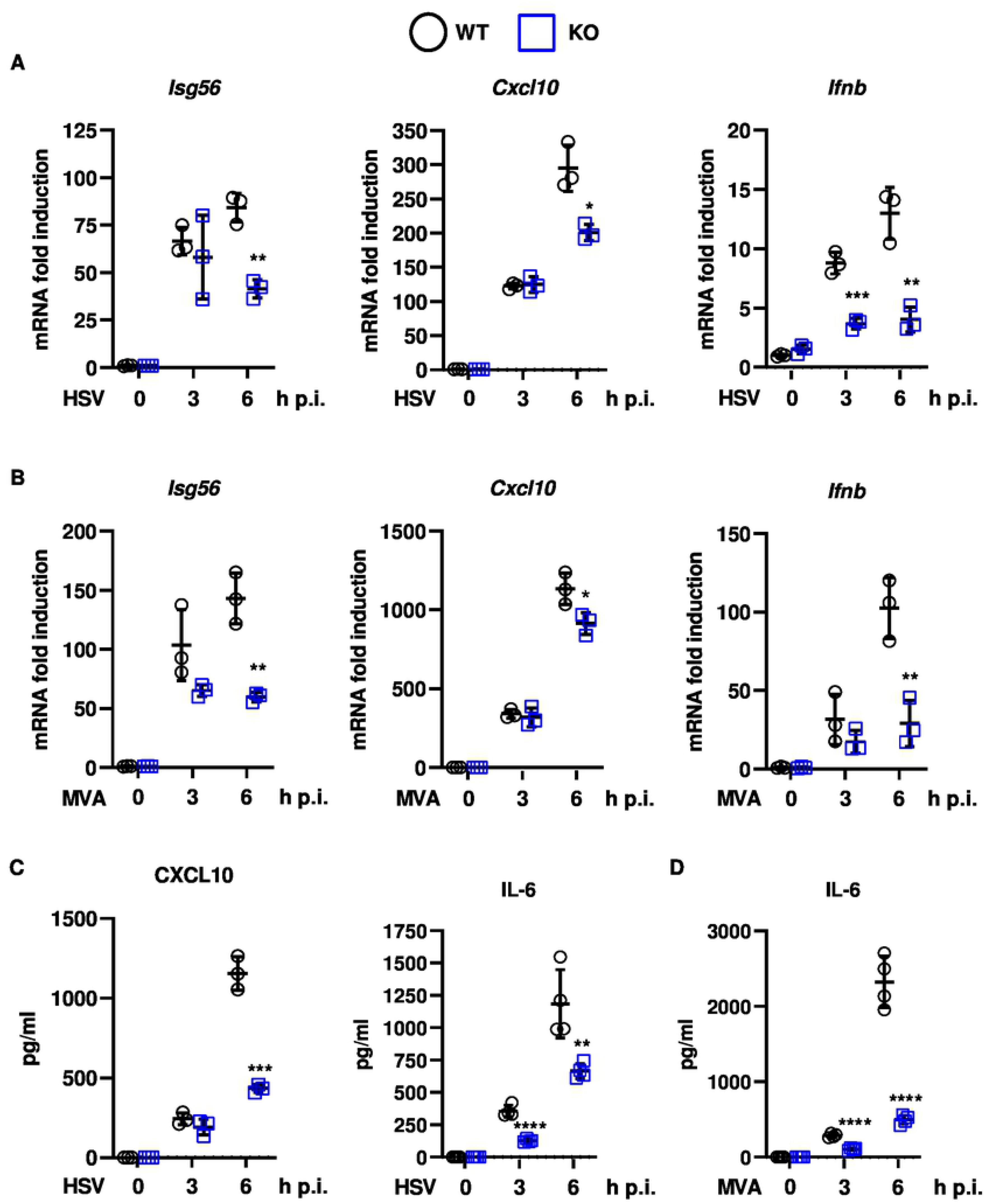
DDX50 is required for IRF3-dependent signalling in response to the dsDNA viruses HSV-1 and VACV. WT MEFs or *Ddx50*^*−/−*^ MEFs were infected for 3 or 6 h at 10 p.f.u./cell with HSV-1 S17 ΔICPO **(A and C)** or MVA **(B and D)** or left uninfected. **(A-B)** mRNA was extracted and *ifnb*, *Isg56* and *Cxcl10* levels were analysed by RT-qPCR relative to *Gapdh*. Representative of at least two independent experiments. **(C-D)** Secretion of CXCL10 and IL-6 were measured at 3 and 6 h post infection by ELISA. Representative of three independent experiments performed in quadruplicate. Statistical significance shown for WT infected vs KO infected.

### Loss of *Ddx50* does not alter IL-1α or TNFα-mediated NF-κB activation

Deletion of *Ddx50* impaired the induction of NF-κB/IRF3-co-transcribed genes in response to dsRNA transfection, ssRNA virus infection (Fig. 1) and dsDNA virus infection (Fig. 2) and previously was reported to modulate MAP kinase signalling (22). Therefore, alternative pathways were tested to determine if the observed defect was specific to RLR signalling. WT or KO MEFs were treated with IL-1α or TNFα and activation of the NF-κB promoter or expression of *Il-6* and *Nfkbia* were measured by Luciferase reporter gene assay or RT-qPCR, respectively. No differences in NF-κB promoter activity or NF-κB-dependent gene expression were observed (Fig. 4A-C), indicating that DDX50 does not play a role in canonical IL-1 receptor- or TNF receptor-induced NF-κB signalling. Collectively, these data suggest DDX50 acts at the stage of IRF3 activation specifically or at or upstream of MAVS and/or TRIF activation before the RNA sensing pathways diverge to activate IRF3 and NF-κB.

**Fig. 4.**
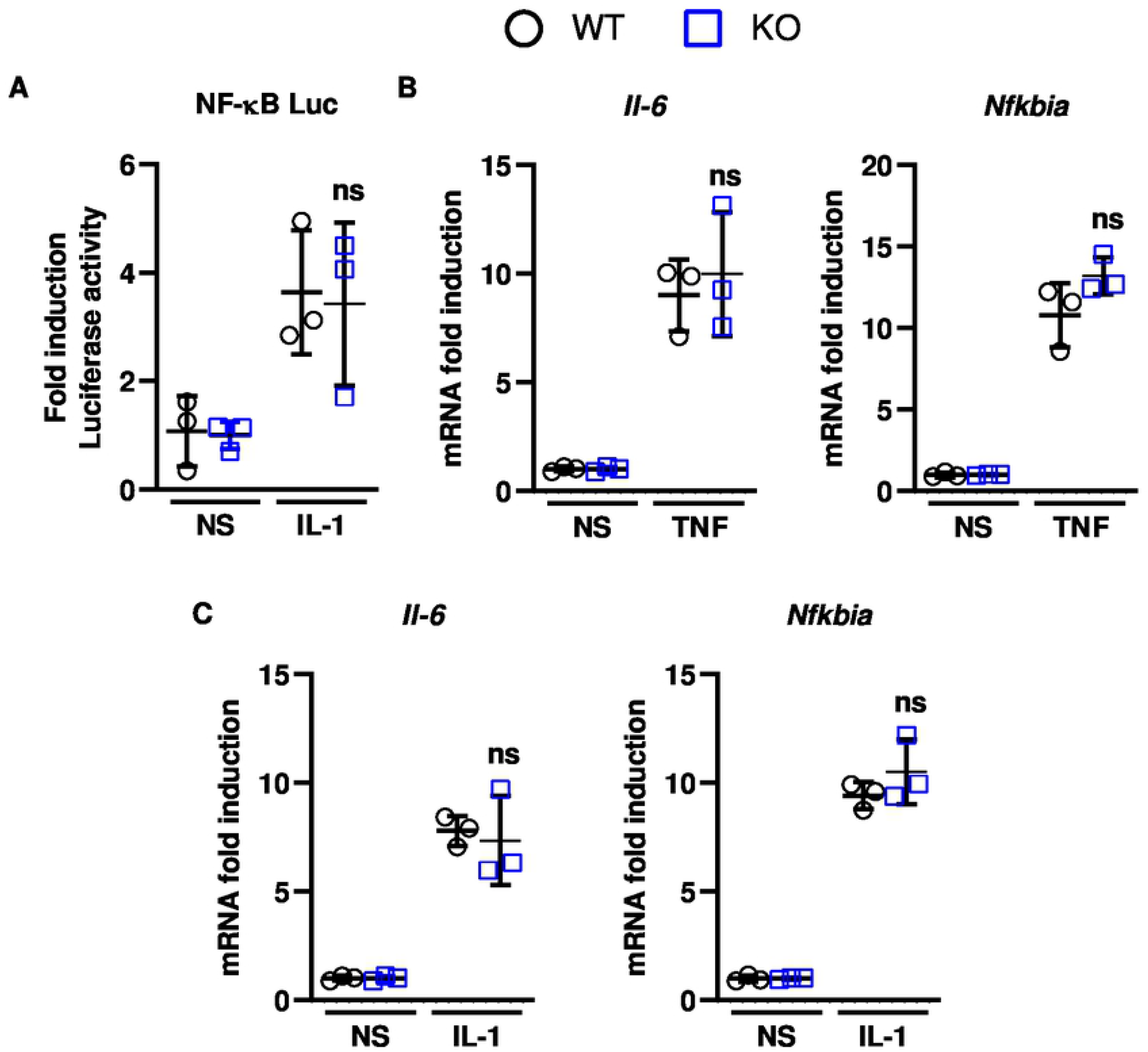
DDX50 (RH-II/Guβ) is not required for NF-κB-dependent gene transcription. **(A)** WT or *Ddx50*^*−/−*^ MEFs were transfected with pNF-κB-Luc or pTK-RL, as an internal control. Cells were left untreated or stimulated for 7 h with 100 ng/ml IL-1α and Firefly Luciferase activity was measured. **(B and C)** WT or *Ddx50*^*−/−*^ MEFs were left untreated or stimulated for 1 h with 100 ng/ml TNFα **(B)** or IL-1α **(C)**. Following mRNA extraction, the fold induction of *Nfkbia* and *Il-6* mRNA levels relative to *Gapdh* were analysed by RT-qPCR. Representative of 3 independent experiments. Statistical significance shown for WT infected vs KO infected.

### DDX50 accumulates in the cytoplasm to activate signalling upstream to MAVS

To investigate where DDX50 acts in the pathway and determine how it facilitates activation of IRF3/NF-κB in response to dsRNA, the phosphorylation of IRF3 was examined. This is a key step in IRF3-dependent signalling and leads to IRF3 dimerisation, nuclear translocation and IRF3-dependent gene transcription. Interestingly, DDX50 was observed to be important for IRF3 phosphorylation at Ser386/396 following PolyIC transfection of both MEFs and HEK293Ts (Fig. 5A-B), mapping DDX50 function to upstream of IRF3 phosphorylation. DDX50 is reported to reside in the nucleolus. However, to act upstream of IRF3 phosphorylation in the canonical cascade one would expect DDX50 to be cytosolic. To explore this further biochemical fractionation of MEFs and anti-DDX50 immunoblotting with or without prior pathway stimulation was used to assess the subcellular localisation of DDX50 in response to cytoplasmic dsRNA. LaminA/C and α-tubulin served as nuclear and cytoplasmic fraction controls, respectively. As described, under resting conditions, the majority of DDX50 was in the nuclear fraction (21) (Fig. 5C). However, DDX50 accumulated in the cytoplasm 1 h post-stimulation (Fig. 5C). At 2 h post-stimulation the level of DDX50 in the cytoplasm returned to basal levels (Fig. 5C). To support this finding the assay was repeated and the localisation of HA-tagged DDX50 was analysed by immunofluorescence. Under resting conditions DDX50 was restricted to the nucleolus, with weak nuclear staining. In agreement with the biochemical fractionation assay, accumulation of DDX50 in distinct cytoplasmic puncta was observed 1 h post infection with SeV (Fig. 5D).

**Fig. 5.**
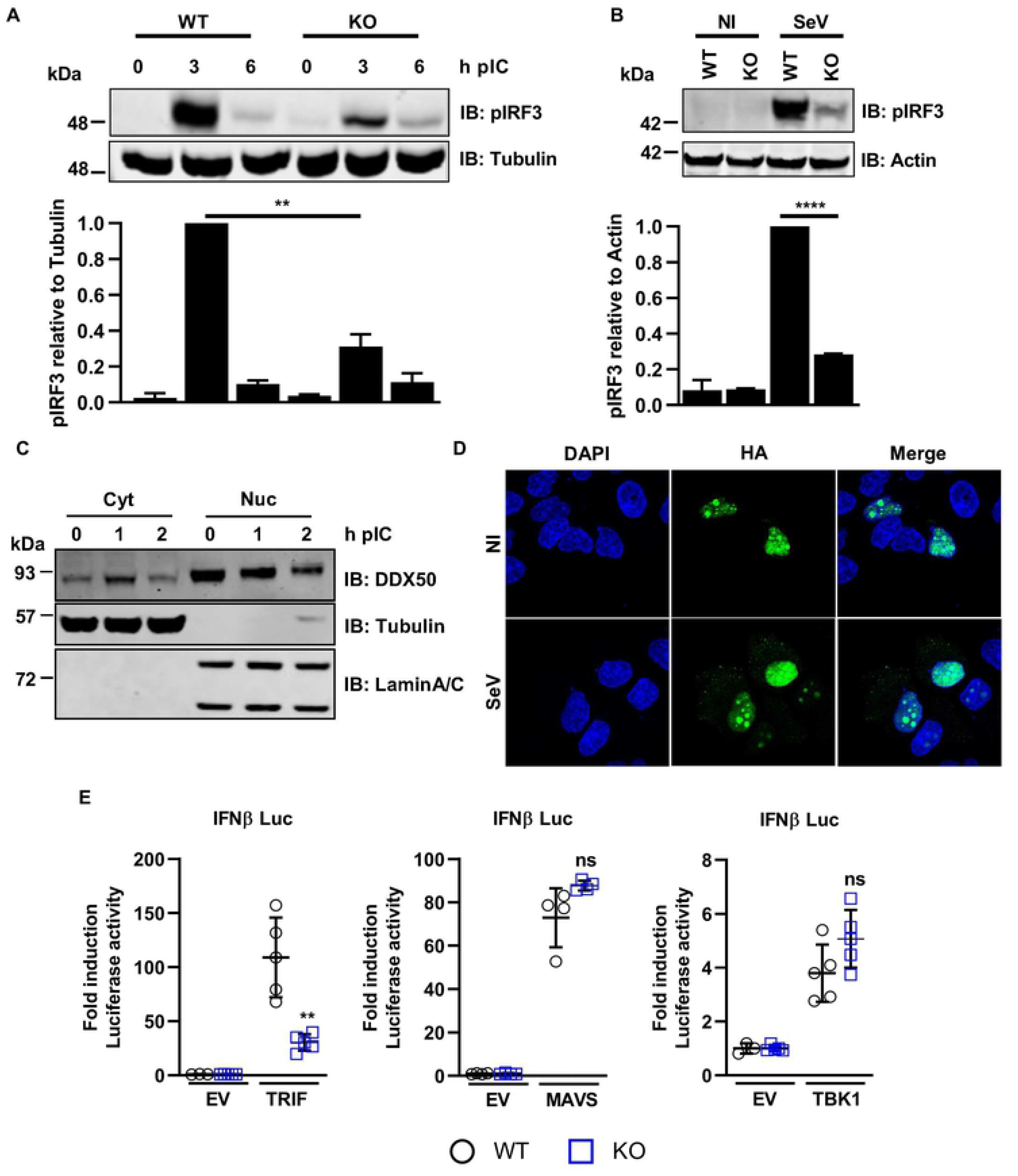
DDX50 accumulates in the cytoplasm in response to cytoplasmic dsRNA and acts upstream to, or independent of, MAVS activation. **(A and B)** Representative immunoblot of phosphorylated IRF3 at **(A)** Ser396 or **(B)** Ser386 (pIRF3) for **(A)** WT or *Ddx50*^*−/−*^ MEFs transfected with lipofectamine only or 5 μg/ml PolyIC for 3 and 6 h or **(B)** WT and DDX50^−/−^ HEK293Ts untreated or infected with SeV for 18 h. Level of IRF3 phosphorylation was calculated by densitometry, relative to α-tubulin **(A)** or actin **(B)** and is representative of at least two independent experiments. **(C)** Representative immunoblot following transfection of WT MEFs with 2.5 μg/ml PolyIC for the indicated times and isolation of the cytoplasmic (cyt) and nuclear fractions (nuc). Immunoblots were stained for DDX50 or α-tubulin and laminA/C as cytoplasmic and nuclear fraction controls, respectively. Representative of three independent experiments. **(D)** Immunofluorescence staining for DDX50 localisation. HeLa cells were transfected with pLDT-DDX50-HA and left uninfected (NI) or infected for 1.5 h with SeV. DDX50 localisation was visualised using an anti-HA antibody. DAPI was used to stain the nucleus. Representative of three independent experiments. (**E**) Luciferase activity of WT or *Ddx50*^*−/−*^ MEFs co-transfected with EV or indicated plasmids along with plasmids encoding Firefly Luciferase under the *Ifnβ* promoter and Renilla as an internal control. Experiments shown are representative of at least three independent experiments. Statistical significance shown for WT stimulated vs KO stimulated.

The nucleocytoplasmic shuttling of DDX50 upon stimulation led us to investigate at which stage in the activation of the IRF3/NF-κB pathway DDX50 might function. The IRF3/NF-κB pathway can be activated by transfection and overexpression of key proteins acting at specific stages of the pathway. Therefore, to map in more detail where DDX50 acts, plasmids encoding TBK1, MAVS or TRIF were co-transfected into WT or KO MEFs along with pLuc-IFNβ and pTK-RL. Activation of the pathway was measured by Firefly and Renilla Luciferase activation as before. No differences in fold activation were observed between the WT and KO cells upon expression of TBK1 or MAVS (Fig. 5E). However, activation was significantly impaired in the KO cell line upon expression of TRIF, mapping DDX50 upstream to or independent of MAVS, but at or downstream of TRIF activation. Notably, DDX50 shares 55.6 % amino acid identity with DDX21, which is essential for TRIF recruitment to MAVS via complex formation with DDX1 and DHX36 in response to cytoplasmic dsRNA (15).

### DDX50 augments TRIF recruitment to activate signal transduction

An essential TRIF-binding domain of DDX21 has been mapped to residues 467-487 within the RNA helicase C domain (15). Strikingly, this motif shares 86% amino acid identity with DDX50 (Fig. 6A). This level of homology was specific for DDX50 and not due to the helicase C domain consensus sequence, because it was not detected within other DExD-box family members such as DHX36 (Fig. 6A). Due to the high level of identity between DDX21 and DDX50 we investigated whether DDX50 can co-immunoprecipitate the DDX1-DDX21-DHX36-TRIF complex. To this end, co-immunoprecipitation assays were performed using extracts of MEF cell lines that stably expressed DDX50-HA and that were transfected with TRIF-cTAP or GFP-Flag. Following stimulation, DDX50-HA specifically co-immunoprecipitated TRIF-cTAP (Fig. 6B). This was confirmed by reciprocal immunoprecipitation in HeLa cells, where hDDX50-HA specifically co-immunoprecipitated with TRIF-cTAP (Fig. 6C). Due to the quality of available anti-TRIF antibodies, co-immunoprecipitation of endogenous TRIF could not be tested. However, endogenous DDX1 did co-immunoprecipitate with DDX50-HA, albeit at low levels (Fig. 6D). This led to the hypothesis that DDX50 may form a cytoplasmic RNA sensing complex with TRIF, to activate TRIF-dependent NF-κB and IRF3 activation. To test this hypothesis, the ability of TRIF to form a complex with DDX1 in WT or DDX50 KO cells was investigated. Interestingly, in the absence of DDX50 the co-immunoprecipitation of endogenous DDX1 with TRIF was diminished, indicating DDX50 may facilitate optimal TRIF recruitment to the RIG-I/MDA5 independent RNA sensing complex (Fig. 6E).

**Fig. 6.**
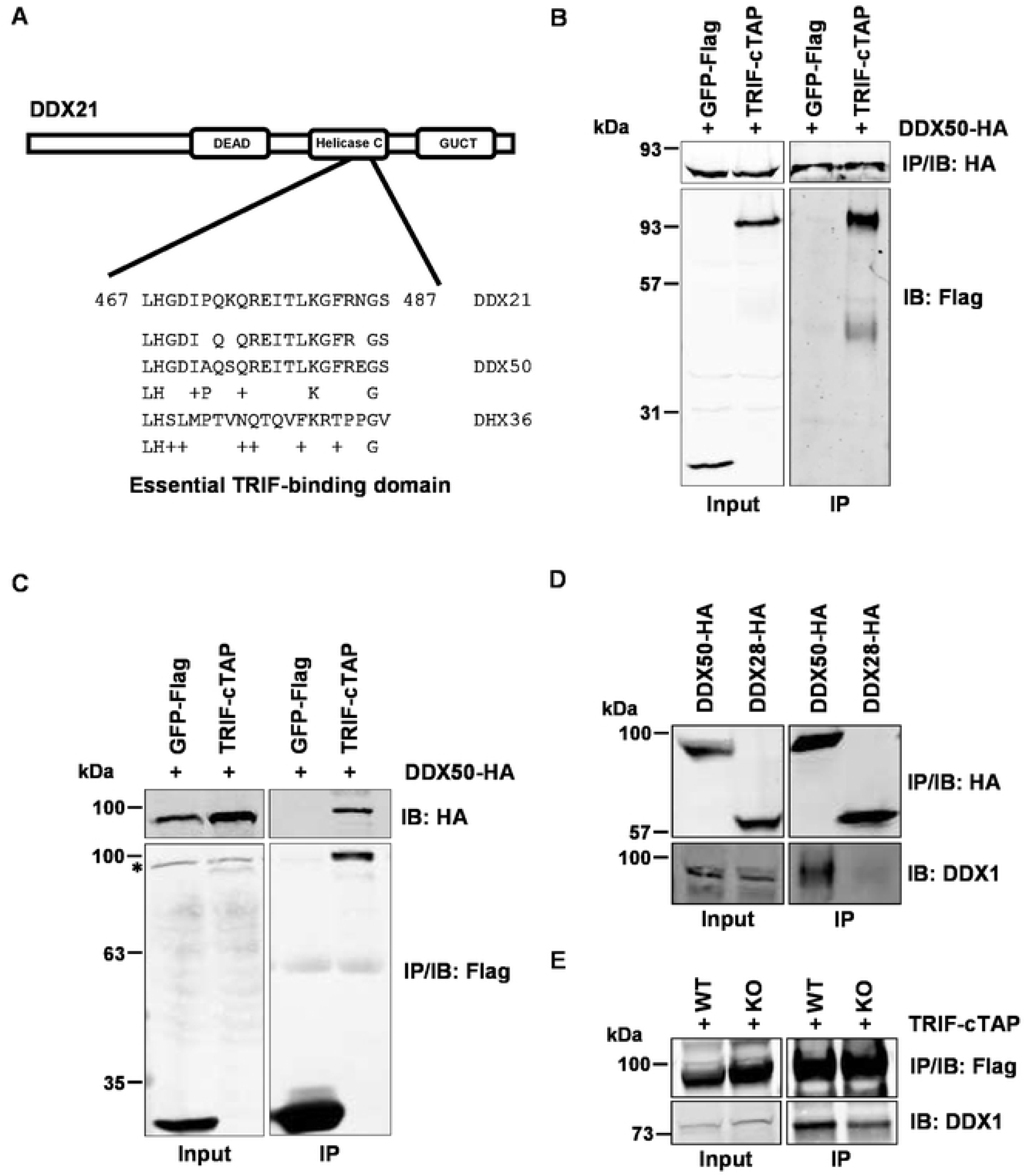
DDX50 co-IPs TRIF and DDX1 and facilitates complex formation. **(A)** Schematic depicting the essential TRIF-binding domain of human DDX21 and the corresponding homologous region in DDX50 or DDX1 and DHX36 as a comparison. **(B)** Immunoblots from co-IP experiments of MEF DDX50-HA cell lines transiently transfected with GFP-Flag and TRIF-cTAP. **(C)** Immunoblots from co-IP experiments of HeLa cell lines transiently transfected with DDX50-HA along with GFP-Flag or TRIF-cTAP. Representative of two independent experiments. *, non-specific band. **(D)** Immunoblots from co-IP experiments of HeLa cell lines transiently transfected with DDX50-HA or DDX28-HA and blotting for endogenous DDX1. **(E)** Immunoblots from co-IP experiments of HEK293T WT or DDX50 KO cells transiently transfected with TRIF-cTAP and blotting for endogenous DDX1. Experiments are representative of three independent experiments. IB, immunoblot; IP, immunoprecipitation.

### DDX50 is a viral restriction factor

IRF3 is a crucial viral restriction factor that controls the transcriptional upregulation of cytokines, chemokines, viral restriction factors and type I IFNs and thereafter IFN-stimulated genes (ISGs) downstream of IFN-induced signalling. Given the role of DDX50 in IRF3/NF-κB-dependent type I IFN production and the synthesis of cytokines during viral infection, its potential as a viral restriction factor was investigated. WT and DDX50 KO cells were infected at either high MOI or low MOI with the dsDNA viruses VACV (MEF, MOI 5 or 0.0001; HEK293T, MOI 5 or 0.0003) and HSV-1 (MOI 0.01) or ZIKV (MOI 1 or 0.1), a ssRNA virus, and virus replication and dissemination were analysed by virus titration and plaque formation. VACV infection produces both single enveloped intracellular mature virus (IMV) and double enveloped cell associate enveloped virus (CEV) and extracellular enveloped virus (EEV) (25). CEVs induce the formation of actin tails to propel virions towards uninfected neighbouring cells. Alternatively, EEVs are released from infected cells and mediate long range dissemination (26). To investigate if loss of *Ddx50* alters viral replication or release, VACV strain Western Reserve (WR) encoding GFP fused to the virus capsid protein A5 (A5-GFP VACV) was used to infect WT or KO MEFs/HEK293Ts at MOI 5 and the total virus or extracellular virus titres 24 h p.i. were determined by plaque assay. No differences in titres of cell associated virus (IMV plus CEV) or released virus (EEV) were observed (Fig. 7A-B) and equal amounts of EEV were produced (approximately 2 % of the total titre; Fig. 7A). Next, virus dissemination and replication were assessed at low MOI. Monolayers of WT or KO MEFs/HEK293Ts were infected at MOI 0.0001 or 0.0003 with A5-GFP-VACV or at MOI 0.01 with HSV-1 strain 17 (S17) encoding GFP fused to Vp26 (Vp26-GFP) and viral titres were determined. Loss of DDX50 in MEFs and HEKs conferred an approximate 6- and 3.5-fold increase in the yield of VACV at 24 and 48 h p.i., respectively (Fig. 8A-B). This difference was not restricted to VACV, and loss of DDX50 resulted in an increase in yield of HSV-1 following low MOI (Fig. 8C). In line with higher viral titres, synthesis of the VACV specific late gene product D8 was enhanced in KO MEFs (Fig. 8F). Notably, the number of plaques formed by VACV was increased on the KO MEFs and HEK293Ts compared to control cells (Fig. 8E and G; Supp Fig. S3A). This suggests that DDX50 restricts plaque formation when cells are infected at low MOI and without DDX50 a greater proportion of virus particles entering cells escape host defences and establish a plaque. Consistent with this observation, complementation of KO MEFs with pCW57-*Ddx50*-HA but not the empty vector (EV) reduced viral yields and plaque formation efficiency to WT levels (Fig. 8G-H). Furthermore, overexpression of hDDX50-HA but not hDDX28-HA in WT human fibroblasts restricted VACV, resulting in significantly lower viral titres (Fig. S3B).

**Fig. 7.**
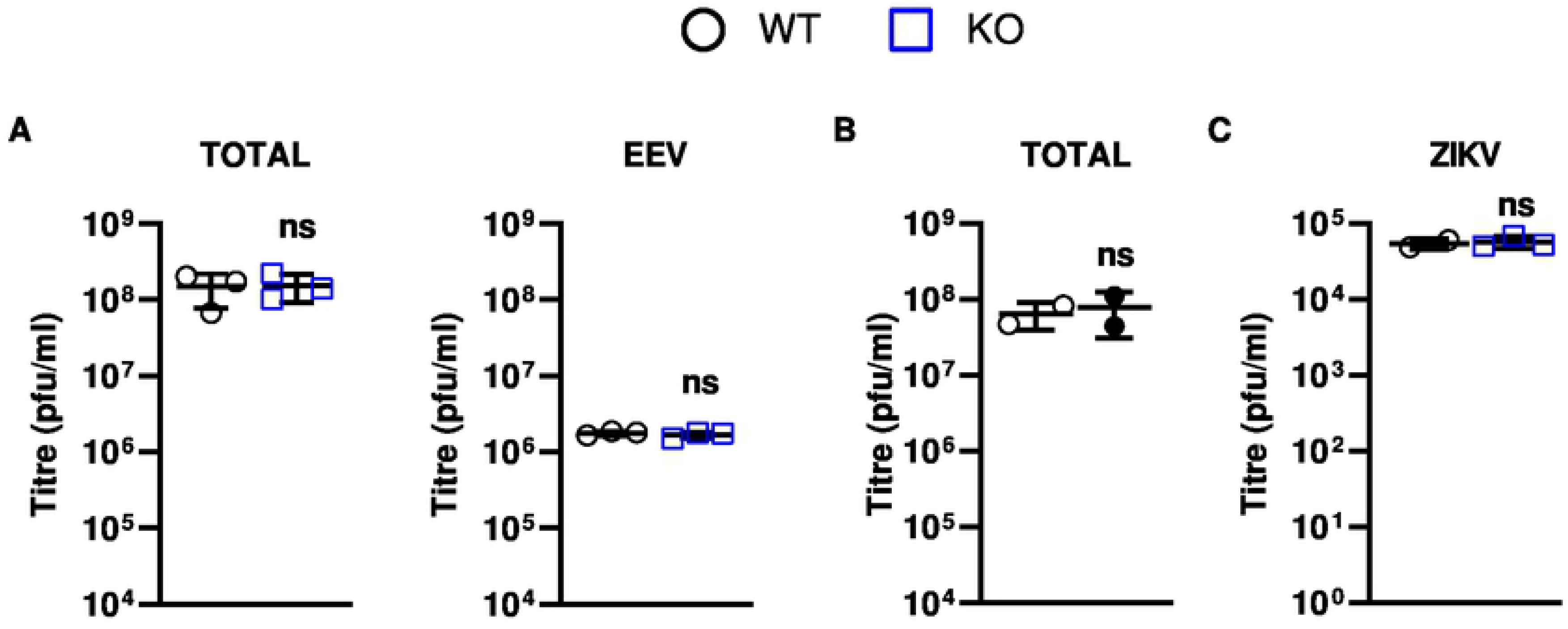
DDX50 does not impact viral replication after high MOI. **(A)** WT or *Ddx50*^*−/−*^ MEFs were infected with A5-GFP VACV WR at 5 p.f.u./cell for 24 h. Viral titres were determined by plaque assay of the medium only (EEV) or total (medium plus cells) on BSC-1 cells. Average of three independent experiments. **(B and C)** WT or *Ddx50*^*−/−*^ HEK293Ts were infected with **(B)** A5-GFP VACV WR at 5 p.f.u./cell for 16 h or **(C)** ZIKV at MOI 1 for 72 h. Viral titres were calculated by titration of cell lysates on Vero E6. Average of two independent experiments. Statistical significance shown for WT vs KO.

**Fig. 8.**
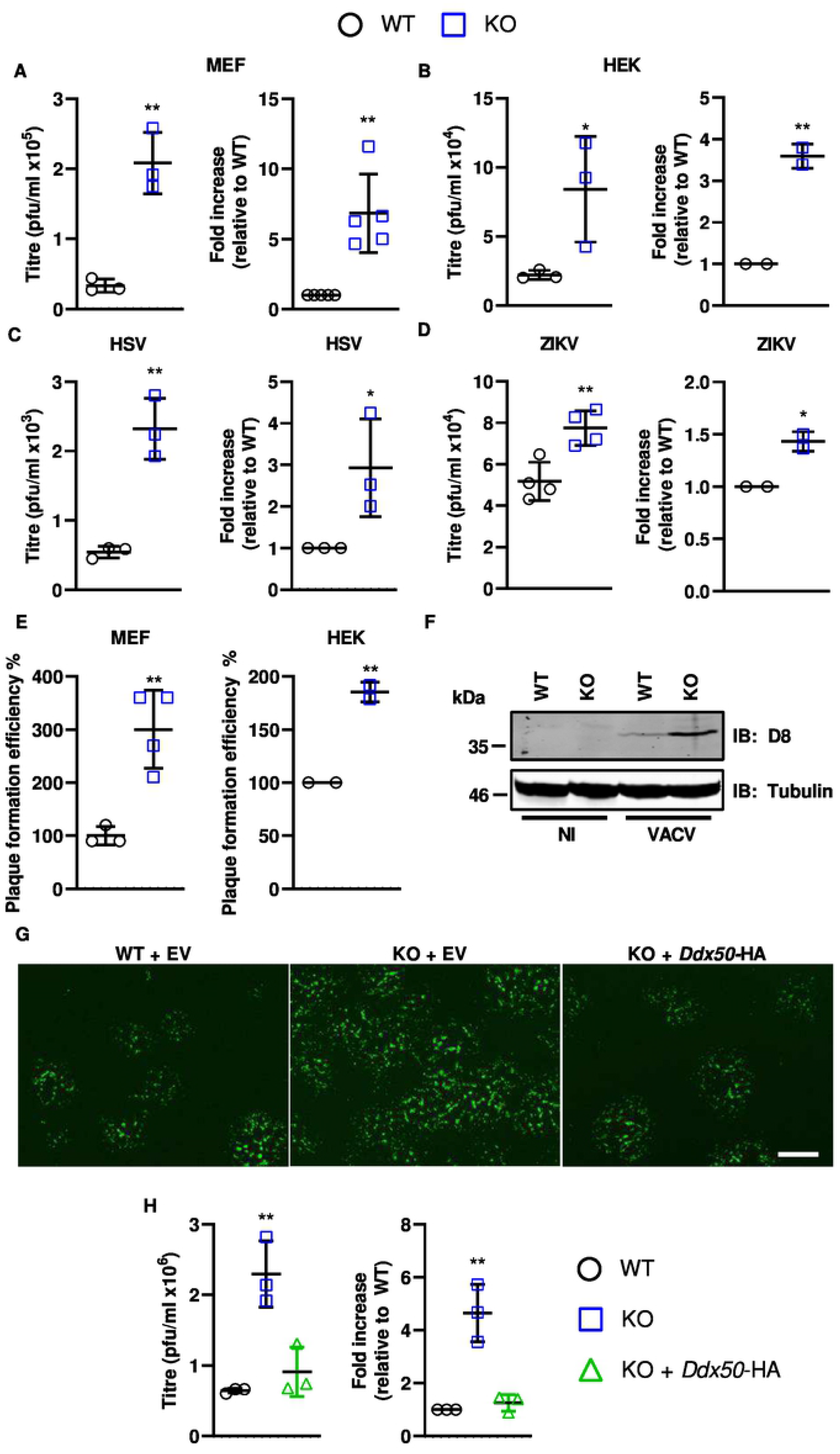
DDX50 is a viral restriction factor. **(A-B)** Monolayers of WT or *Ddx50*^*−/−*^ MEFs were infected with A5-GFP VACV WR at 80 p.f.u. (MOI = 0.0001). Viral titres at 24 h p.i. were determined by plaque assay on BS-C-1 cells and are represented as p.f.u./ml (left panel) or fold increase in replication relative to WT cells (right panel). **(B)** As in **(A)** but using WT or *Ddx50*^*−/−*^ HEK293Ts infected at MOI 0.0003 for 48 h. Results shown are representative of 2 independent experiments. **(C)** Monolayers of WT or *Ddx50*^*−/−*^ MEFs were infected with HSV-1 S17 Vp26-GFP at 6600 p.f.u. (MOI = 0.01). Viral titres at 48 h p.i. were determined by plaque assay on U2OS cells and are represented as p.f.u./ml (left panel) and fold increase in replication relative to WT cells (right panel). **(D)** WT or *Ddx50*^*−/−*^ HEK293Ts were infected with ZIKV at MOI 0.1 for 72 h. Titres were determined by plaque assay on Vero E6 cells and data are shown as for A-C. Titres shown are an average of 2 independent experiments. **(E)** Cells were infected at low MOI as in **(A)** and **(D)** and plaque numbers were enumerated 24 h p.i.. Data are expressed as the plaque formation efficiency on KO cells compared to WT cells. Representative of two independent experiments**. (F)** WT or *Ddx50*^*−/−*^ MEFs were infected with A5-GFP VACV WR at MOI = 0.0001 and expression of the VACV late protein D8 was analysed by immunoblot at 24 h p.i. Representative of two independent experiments. **(G-H)** WT or *Ddx50*^*−/−*^ MEFs transduced with pCW57-EV or pCW57-*Ddx50*-HA were infected as in **(G)**. Representative fluorescence images of plaque morphology following infection. Scale bar, 500 μM. **(H)** Viral titres at 24 h p.i. from cells infected as in **(G)** For all panels unless stated otherwise titres shown are representative of at least 3 independent experiments and fold changes shown are an average of at least 2 independent experiments. Statistical significance is shown for WT EV vs KO EV.

Given that the activation of IRF3 restricts RNA virus infection as well, a ZIKV replication assay was performed in the absence of DDX50. Parental HEK293T and derived DDX50−/− cells were infected with ZIKV at MOI 1 or 0.1. Three days p.i., supernatants of infected cells were collected and infectious virus was titrated by plaque assay on Vero E6 cells. The absence of DDX50 increased ZIKV replication following low MOI (Fig. 8D), however in concordance with dsDNA viral infection, no difference was observed at high MOI (Fig. 7C). This suggests that the role of DDX50 in promoting activation of the IRF3 pathway contributes to the restriction on ZIKV infection. Together, these results provide evidence that DDX50 promotes antiviral signalling during infection and is a restriction factor for both DNA and RNA viruses, with its loss resulting in increased viral spread and subsequent replication in tissue culture.

## Discussion

Type I IFNs are critical regulators of antiviral immunity and infection control and therefore, understanding the mechanisms leading to their production during infection is important. During the last decade, much research has studied the canonical RLRs and RNA sensors RIG-I, MDA5 and TLR3, and has investigated their activation, expression and mechanisms of regulation in response to RLR agonists and infection. Zhang and colleagues described a TLR3, RIG-I and MDA5-independent pathway in mouse dendritic cells in which cytoplasmic RNA was sensed by a complex consisting of DDX1-DDX21-DHX36, leading to recruitment of TRIF (15). Here, DDX50 is described as a new component of the RNA sensing signalling pathway. DDX50 is a TRIF-binding RNA helicase that is an integral member of TRIF-dependent IRF3/NF-κB activation in fibroblast/epithelial cells. Aside from the initial *in vitro* characterisation of the RNA helicase functional domains of DDX50, little is understood about its cellular role. A previous study concluded that DDX50 is required for MAP kinase activation through c-Jun binding (22). However, whilst a defect in RNA sensing and signalling was observed here, no differences in TNFR/IL-1R-dependent NF-κB signalling were detected. Differences in the signalling cascade that were observed are independent of MAP kinase and activator protein 1 (AP-1) activation, indicating that this is an independent role for DDX50.

DDX50 is needed for optimal IRF3/NF-κB-dependent gene expression, and cytokine synthesis and secretion following stimulation with dsRNA analogue PolyIC, SeV infection or infection with the dsDNA viruses HSV-1 and VACV. Further investigation found that without DDX50, IRF3 phosphorylation was impaired downstream of these stimuli, but that signalling was intact following activation via MAVS overexpression. This mapped the activity of DDX50, a nucleolar protein, to early in the signalling cascade upstream or independent of MAVS activation. DDX50 shuttling and cytoplasmic accumulation in distinct puncta upon stimulation is reminiscent of DDX1/TRIF staining in response to PolyIC treatment and is consistent with a role for DDX50 in cytoplasmic regulation of IRF3 signalling (15). The RNA sensing complex consisting of DDX1, DHX36 and DDX21 identified by Zhang and colleagues did not report on DDX50. However, this was performed in mouse dendritic cells and the expression of DDX50 varies from cell type to cell type (human atlas data). It may be that DDX50 plays a more significant role in non-haematopoietic cells or that it was below detection in the initial screen. Alternatively, the high sequence identity between DDX50 and DDX21 may have masked the role of DDX50 following knockdown of DDX21. In the DDX1-DHX36-DDX21 complex, DDX21 acts as a scaffold to recruit TRIF upon DDX1 agonist binding. Therefore, we hypothesise that DDX50 may act in a similar fashion. Notably, whilst DDX21 and DDX50 have non-redundant roles, DDX50 is essential for DDX21 helicase activity *in vitro* (21). So, it is possible that DDX50 may function to support DDX21 or even DDX1 (the RNA binding protein) activity in this complex, which may explain why DDX50 is not functionally redundant. Whilst diminished binding of TRIF and DDX1 was observed in the absence of DDX50, it was not abolished, and the role of the functional domains of DDX50, DDX21 and DDX1 in RNA sensing warrants future experimentation.

Consistent with a role for DDX50 in innate immune signalling, DDX50 is shown to be a viral restriction factor. Loss of DDX50 resulted in an attenuated immune response to infection with VACV or HSV-1 and enhanced replication of VACV, ZIKV and HSV-1 in tissue culture after low MOI infection. Notably, a greater number of VACV plaques were formed on KO cell lines suggesting that DDX50 acts to restrict viral infection and in its absence a greater proportion of infecting virus particles escape host defences and lead to plaque formation. At high MOI, there were no differences in virus yield suggesting that infection of a single cell by many incoming virus particles can overcome DDX50-mediated restriction. This is reminiscent of cellular restriction factors involved in innate immune signalling. These data provide evidence of the biological relevance of DDX50 for antiviral signalling during infection. Interestingly, DDX50 is reported to co-immunoprecipitate with the positive sense RNA virus Dengue (DENV) RNA (27) and recent publications using siRNA to knockdown DDX50, suggest that it may inhibit DENV replication (28). Following knockdown, the authors reported a reduction in IFNβ promoter activity and therefore hypothesised that DDX50 may regulate type I IFN production during DENV infection (29). This is consistent with our findings that the ZIKV titre is increased in the absence of DDX50, and together provides evidence that DDX50 is a viral restriction factor in response to multiple RNA and DNA viruses. Therefore, DDX50 as a restriction factor may extend beyond the viruses tested in this study and act broadly to detect viral RNA and restrict viral replication through activation of IRF3-dependent gene transcription.

Although DDX50 was required for optimal signal transduction its absence did not abolish signalling in response to viral infection or stimulation. Given that DDX50 binds TRIF, a protein that is non-essential for RIG-I/MDA5 signalling, and that DDX1 acts independent of canonical RIG-I signalling, we propose that DDX50 acts in concert with other receptors for optimal antiviral signalling and restriction. Whilst this study identifies a role for DDX50 in RNA sensing, both HSV-1 and VACV are DNA viruses. HSV-1 is reported to be restricted mostly by the cGAS-STING pathway (30). However, DNA sensing and antiviral signalling is positively regulated by both TRIF and RNA sensing during HSV-1 infection (31–33), highlighting the essential role of RNA sensors during DNA virus infection. Cells infected with VACV contain large amounts of dsRNA late during infection (34,35). This is due to the virus intermediate and late genes lacking specific transcriptional termination sequences and so lengthy overlapping transcripts are produced that hybridise to form dsRNA (36). These transcripts can be sensed and activate innate immune signalling pathways (37). In addition, such dsRNA can bind to and activate IFN-induced proteins such as PKR and 2’-5’ oligoadenylate synthetase (OAS) to mediate translational shutoff. The importance of dsRNA in activating host defences is illustrated by the fact that VACV, despite being a dsDNA virus, encodes a dsRNA binding protein called E3 (38), that contributes to virulence (39). It is important to note that TRIF is also an essential component of the STING pathway (31). Therefore, the level to which DDX50 restricts DNA viruses in an RNA-sensing dependent manner, or whether it can further influence TRIF signalling in the cGAS-STING pathway, warrants future investigation. Furthermore, the importance of DDX50 in RNA sensing and its contribution in antiviral immunity requires validation *in vivo*. Unfortunately, to date there are no KO mice or models available, however with the recent success in generating *Ddx21* KO mice, it may soon be a plausible avenue for investigation.

In conclusion, the DExD-Box RNA helicase DDX50 is identified as a crucial component in the host cell RNA sensing machinery, acting to facilitate IRF3 activation and inhibit viral dissemination. It is proposed that DDX50 may act through the recruitment of TRIF to the DDX1 RNA sensing complex.

## Acknowledgements

We would like to thank Dr. B.J. Ferguson, University of Cambridge and Dr. A. Shenoy, Imperial College London, for their helpful feedback and advice and Dr. Trevor Sweeney, University of Cambridge for providing ZIKV for this project. pCW57-GFP-2A-MCS was a gift from Adam Karpf (Addgene plasmid #71783).

## Material and Methods

### Cells, plasmids, reagents and viruses

All reagents were purchased from Sigma unless stated otherwise. BSC-1 (ATCC CCL-26), U20S (ATCC HTB-96), HEK293T (ATCC CRL-11268) and immortalised mouse embryonic fibroblasts (MEF) were all grown in Dulbecco’s modified Eagle’s medium (DMEM) high glucose (Gibco), supplemented with 10 % foetal bovine serum (FBS; Pan Biotech), 50 μg/ml penicillin/streptomycin (P/S), non-essential amino acids (NEAA). HeLa (ATCC CCL-2) and human fibroblasts (HF) clone EF-1-F (sourced from Doorbar lab, University of Cambridge) were grown in MEM (Gibco) supplemented with 10 % FBS, 50 μg/ml P/S and non-essential amino acids (NEAA). All cells were grown at 37 °C in a 5 % CO_2_ atmosphere and were routinely screened for mycoplasma contamination. All plasmids constructed in this study are listed in Table S1. Vaccinia virus (VACV) strain Western Reserve (WR) recombinant vA5-GFP (40), modified vaccinia virus Ankara (MVA) (41), HSV-1 S17 GFP-Vp26 (42) and HSV-1 ΔICPO (43) were described. The titre of infectious viral particles (plaque-forming units per ml, p.f.u/ml) was determined by plaque assay on BSC-1 cells for VACV WR and on U2OS for HSV-1. Sendai virus Cantell strain (Licence No. ITIMP17.0612A) was a gift from Steve Goodbourn, St George’s Hospital Medical School, University of London. ZIKV engineered to express a mCherry marker (44) was a kind gift from Dr. Trevor Sweeney, Department of Pathology, University of Cambridge.

### CRISPR-cas9 generation of knockout cell lines

Guide RNA design and synthesis, and pX459 plasmid construction was performed following the Zhang lab protocol (45). Specific guide RNAs are described in Table S1. To generate KOs, MEFs were transfected with pX459 plasmids using LT1 following the manufacturer’s protocol. Twenty-four h post transfection MEFs and HEK293Ts were treated with 4 μg/ml and 1 μg/ml puromycin (Invitrogen) for 48 h, respectively. Single cell clones were selected by limiting dilution, expanded, and screened for DDX50 protein levels by immunoblot. To confirm successful knockouts, the genomic DNA of selected clones was purified following the manufacturer’s protocol (Qiagen, QIAamp DNA mini kit). *Ddx50* was amplified using the primer pair gagcgtccttcctggagattg / ctcaagtctgcccatctctcg and *DDX50* was amplified using the primer pair ctgtgtcaccaggtggcatg / gactcgtgtaactttctttccc. Single allele PCR amplicons were then cloned into pCR2.1-TOPO by blunt end ligation (Thermofisher) and 10 clones were sequenced for each KO cell line clone. Single allele sequencing results were compared to the sequence results of the gDNA PCR amplicon to check all alleles had been identified and that all mutations resulted in frameshift truncations.

### pLDT and pCW57 cell line generation

WT and *Ddx50*^*−/−*^ MEF and WT HF cell lines inducibly overexpressing DDX50 were obtained by transduction using lentivirus vectors. pLDT and pCW57 cell lines were generated as described (46) with the following alterations. MEFs and HFs were selected in 4 μg/ml puromycin (Invitrogen), followed by single cell selection. For HF pLDT-TetR-GFP was co-packaged along with the pLDT-MCS plasmids and selected for with 500 μg/ml neomycin (Gibco).

### Luciferase reporter assay

HEK293T, HF and MEF cell lines were transfected with 10 ng of the internal control plasmid pTK-Renilla (pRL-TK, Promega) or 60 ng of the reporter plasmid pLUC-NF-κB (R. Hofmeister, University of Regensburg, Germany) or pLUC-IFNβ (T. Taniguchi, University of Tokyo) using LT1 transfection reagent and following the manufacturer’s instructions (MirusBio Ltd). Where stated, plasmids encoding TRIF, MAVS or TBK-1 (K.A. Fitzgerald, University of Massachusetts Medical School) were co-transfected. Twenty-four h post-transfection, cells were stimulated with IL-1α (Invivogen) or TNFα (Invivogen) at 100 ng/ml or transfected with 5 μg/ml high molecular weight (HMW) PolyIC (Invivogen) using Liopfectamine 2000 (Invitrogen), or mock-transfected with lipofectamine only, or treated exogenously with 5 μg/ml PolyIC, or left unstimulated for 6 h in DMEM or MEM with 2 % FBS. Alternatively, cells were stimulated by SeV infection at 1:100 dilution of stock for 24 h. Following stimulation cells were lysed in 1 × Passive lysis buffer (Promega) and Firefly luciferase and Renilla luminescence were measured using the MARS data analysis software on the FLUOstar Omega Luminometer (BMG Labtech). Relative luminescence levels were calculated by normalising Firefly luminescence to Renilla and data are presented as relative to the non-stimulated untreated condition, or EV where relevant, for each cell line. Each condition was performed with quadruplicate technical replicates and is representative of two biological repeats.

### ELISAs and RT-qPCR

MEFs were seeded in DMEM with 2 % FBS and HEK293Ts were seeded in DMEM with 10 % FBS. After 18 h cells were mock-transfected or transfected with 5 μg/ml HMW PolyIC (Invivogen) using Lipofectamine 2000 (Thermofisher) for 7 h or infected with SeV (Cantell Strain) for 4.5 or 24 h where stated. The culture medium was cleared by centrifugation at 17, 000 × *g* and stored at −20 °C before analysis by ELISA. The level of human or mouse CXCL10/IP-10 was determined using a DuoSet ELISA kit (R&D Systems) and the level of mouse IL-6 was determined using a DuoSet ELISA kit (R&D systems) following the manufacturer’s instructions. Data were collected and analysed using the MARS data analysis software on the FLUOstar Omega Luminometer (BMG Labtech). Experiments were carried out in triplicate and measured with technical repeats, unless stated otherwise. RNA extraction, cDNA synthesis and RT-qPCR were carried out as described previously using first strand synthesis (Invitrogen) (47). qPCR was performed using the primers indicated in Table S2.

### Immunoprecipitations

HeLa cells were transfected with pLDT-hDDX50-HA and co-transfected with pCDNA3-GFP-Flag or pCDNA3-TRIF-cTAP where stated. For MEFs, DDX50-HA pCW57 cell lines were induced with 2 μg/ml doxycycline 24 h prior to transfection with pCDNA3-GFP-Flag or pCDNA3-TRIF-cTAP. WT and *Ddx50*^*−/−*^ HEK293Ts were transfected with pCDNA3-TRIF-cTAP. Twenty-four h post transfection cells were stimulated by transfection with 5 μg/ml PolyIC or infected with SeV (1:200) where stated. Following stimulation, cells were washed and lysed in 50 mM Tris pH 7.6, 150 mM NaCl, 1 % NP40 (IGEPAL CA-630), 1 mM EDTA, 10 % glycerol and supplemented with protease inhibitor. Proteins were immunoprecipitated as described (48) with M2 Flag-beads or HA-beads. After the final wash, beads were incubated in 4 × sample buffer (Tris 0.5 M pH 6.8, 40 % glycerol, 6 % SDS, 1 % bromophenol blue and 0.8 % β-mercaptoethanol), boiled and analysed by immunoblotting.

### Immunoblotting

Samples were prepared by the addition of 4 × sample buffer, boiled and separated by gel electrophoresis in Tris-glycine SDS (TGS) buffer (20 mM Tris, 192 mM glycine, 1 % (w/v) SDS) and transferred to a nitrocellulose membrane (GE Healthcare) in Tris glycine (TG) buffer (20 mM Tris-HCl pH 8.3, 150 mM glycine) using the Turboblot system (BioRAD). Membranes were blocked in 5 % milk in Tris-buffered saline (10 mM Tris, 150 mM NaCl) pH 7.4 with 0.1% (v/v) Tween-20 (TBS-T) for 1 h before incubating with the primary antibody overnight at 4 °C. Primary antibodies: rabbit monoclonal anti-Flag (F7425), anti-DDX50 (Abcam; ab109515), anti-IRF3 Ser386 (Abcam, ab76493), rabbit polyclonal anti-HA (H6908), mouse monoclonal anti-Flag (F1804), anti-α-tubulin (Millipore; 05-829), anti-DDX50 (Santa cruz, sc-81077), anti-DDX1 (Santa cruz; sc-271438), anti-LaminA/C (Abcam; ab8984), mouse polyclonal anti-IRF-3 S396 (CST; #4947S) or mouse monoclonal anti-D8 clone AB1.1 (49). Membranes were washed 3 times in TBS-T before incubating with secondary antibodies for 1 h. Secondary antibodies were goat anti-rabbit IRDye 800CW (926-68032211; LiCOR) and goat anti-mouse IRDye 608LT (926-68020; LiCOR) or, for immunoprecipitated samples, biotin-anti-mouse light chain followed by streptavidin IRDye 680LT (926-68031; LiCOR) was used. Finally, membranes were washed 3 times in TBS-T, dried and imaged using the LiCOR system and Odyssey software. For protein level comparisons, densitometry was calculated using ImageJ.

### Virus growth analysis

To measure viral spread, confluent monolayers of WT or KO MEFs were infected with 80 p.f.u of vA5-GFP or 200 p.f.u of HSV-1 S17 Vp26-GFP in DMEM with 2 % FBS. Alternatively, for the single step virus replication analysis, cells were infected with 5 p.f.u/cell of vA5-GFP. Plates were rocked regularly at 37 °C for 2 h before incubating at 37 °C for the indicated times. Plaques were imaged using an Axiovert.A1 inverted fluorescence microscope connected to a Zeiss MRc colour camera and processed using Axiovision Rel. 4.8 imaging software. To determine the viral titre, the medium and cells were collected, freeze-thawed three times, sonicated at 2.0 for 20 s three times (for VACV only) and titrated on BSC-1 or U20S for VACV and HSV-1, respectively. For ZIKV infection and titration, 3 × 10^6^ parental HEK293T or DDX50^−/−^ cells were seeded on poly-D-lysine pre-coated 6-well plates. Cells were infected with ZIKV at MOI 0.1 the next day. Three infection p.i., supernatants of the infected cells were collected and virus infectivity was titrated by plaque assay on Vero E6 cells. To titrate ZIKV samples, Vero E6 cells on 6-well plates (90% confluence) were infected for 2 h, the inoculum was removed and cells were incubated in MEM with 1.5% carboxymethyl cellulose for 5 d. Cells were then fixed with 4% paraformaldehyde (PFA) and stained with toluidine blue.

### Cell sub-fractionation

Following stimulation for the times indicated, cells were washed in PBS and fractionated using the NE-PER™ Nuclear and Cytoplasmic Extraction Kit following the manufacturer’s protocol (ThermoFisher).

### Immunofluorescence

Briefly, cells were fixed in 4 % PFA/PBS for 20 min, washed in PBS, quenched in 150 mM NH_4_Cl/PBS for 10 min and permeabilised in 0.1 % Triton X-100/PBS for 10 min, before a final wash and block in 5 % FBS/PBS. Cells were stained by inverted incubation in 5 % FBS/PBS with anti-rabbit HA (dilution 1:100) antibody for 1 h, washed in 5 % FBS/PBS and incubated for a further 30 min with the secondary goat anti-rabbit IgG Alexa-Fluor 488 (Jackson immunoresearch; 111-545-003). Coverslips were mounted in Mowiol (10 % w/v Mowiol4–88 (CalBiochem), 25 % v/v glycerol, 100 mM Tris-HCl pH 8.5, 0.5 μg/ml DAPI (4’,6-diamidino-2-phenylindole, Sigma) and images were acquired using a Zeiss LSM780 confocal laser scanning microscopy system and processed using the Zeiss Zen microscope and Axiovision 4.8 software.

### Statistics

All experiments are presented as technical or biological averages where stated. Data presented are the mean +/− SD. All assays were analysed by unpaired T-test with GraphPad Prism 8 Software where p<0.05 = *, p<0.01 = **, p<0.001 = *** and p<0.0001 = ****.

## Supporting information

**Table S1. Constructs and primers used in the study**

**Table S2. Primers for qPCR**

**Fig. S1. CRISPR-Cas9 mediated knockout of *Ddx50/DDX50* (*RH-II/Guβ*) in fibroblasts.**

**Fig. S2. DDX50 overexpression augments nucleic acid sensing.**

**Fig. S3. Overexpression of DDX50 inhibits VACV dissemination and replication.**

